# The *Vibrio* Type III Secretion System 2 is not restricted to the *Vibrionaceae* and encodes differentially distributed repertoires of effector proteins

**DOI:** 10.1101/2022.08.23.504659

**Authors:** SA Jerez, N Plaza, V Bravo, IM Urrutia, CJ Blondel

## Abstract

*Vibrio parahaemolyticus* is the leading cause of seafood-borne gastroenteritis worldwide. A distinctive feature of the O3:K6 pandemic clone, and its derivatives, is the presence of a second, phylogenetically distinct, Type III Secretion System (T3SS2) encoded within the genomic island VPaI-7. The T3SS2 allows the delivery of effector proteins directly into the cytosol of infected eukaryotic cells to subvert key host cell processes, critical for *V. parahaemolyticus* to colonize and cause disease. Furthermore, the T3SS2 also increases the environmental fitness of *V. parahaemolyticus* in its interaction with bacterivorous protists; hence it has been proposed that it contributed to the global oceanic spread of the pandemic clone. Several reports have identified T3SS2-related genes in *Vibrio* and non-*Vibrio* species, suggesting that the T3SS2 gene cluster is not restricted to the *Vibrionaceae* and can mobilize through horizontal gene transfer events. In this work, we performed a large-scale genomic analysis to determine the phylogenetic distribution of the T3SS2 gene cluster and its repertoire of effector proteins. We identified putative T3SS2 gene clusters in 1130 bacterial genomes from 8 bacterial genera, 5 bacterial families and 47 bacterial species. A hierarchical clustering analysis allowed us to define 6 T3SS2 subgroups (I-VI) with different repertoires of effector proteins, redefining the concepts of T3SS2 core and accessory effector proteins. Finally, we identified a subset of T3SS2 gene clusters (subgroup VI) that lack most T3SS2 effector proteins described to date and provided a list of 10 novel effector candidates for this subgroup through bioinformatic analysis. Collectively, our findings indicate that the T3SS2 extends beyond the *Vibrionaceae* family and suggest that different effector protein repertories could have a differential impact on the pathogenic potential and environmental fitness of each bacteria that have acquired the *Vibrio* T3SS2 gene cluster.

**DATA SUMMARY:**

1. All genome sequences used in this study were downloaded from the National Center for Biotechnology Information (NCBI) RefSeq or GenBank databases (See **Table S1** for accession numbers).
2. Files for the T3SS2 reconstructed phylogenetic tree (Newick tree and MSA fasta file), hierarchical clustering data analysis file from MORPHEUS, **Table S1** with genome accession numbers and all the data of the absence/presence of T3SS2-related components, and **Table S2** with the prediction of novel effector proteins are available as part of the online Supporting Dataset at the Zenodo data repository (https://doi.org/10.5281/zenodo.7016552). The T3SS2 phylogenetic tree can be interactively visualized in https://itol.embl.de/tree/19016190125374711626959067#

## INTRODUCTION

*Vibrio parahaemolyticus* is a Gram-negative bacterium that resides in marine and estuarine environments [1, 2]. In 1996, a pandemic clone of the O3:K6 serotype emerged and spread around the globe, becoming the leading cause of seafood-borne gastroenteritis worldwide [3]. Whole-genome sequencing of the pandemic strain RIMD2210633 revealed the presence of a novel Type III Secretion System (T3SS2) encoded within an 80-kb genomic island on chromosome 2, known as the *V. parahaemolyticus* pathogenicity island 7 (VPaI-7) [4]. While all *V. parahaemolyticus* strains encode a T3SS on chromosome 1 (T3SS1), only *V. parahaemolyticus* strains derived from the pandemic clone harbor the phylogenetically distinct T3SS2 [4].

T3SS are complex nanomachines that enable Gram-negative bacteria to deliver proteins, known as effectors, directly from the bacterial cytosol into the cytosol of eukaryotic cells [5, 6]. The translocation of effectors into target cells enables bacteria to subvert a wide variety of host cell functions [7–9]. In addition, the specific repertoire of proteins delivered to infected cells determines which cellular networks are subverted by the bacterium and ultimately influence the outcome of the infection [5, 6, 10]. In this context, even closely related T3SSs can have different repertoires of effector proteins due to horizontal gene transfer (HGT) and/or genomic rearrangement events [11, 12]. A recent global survey described that T3SS are widely distributed among Gram-negative bacteria, identifying 174 nonredundant T3SS from 109 distinct bacterial genera [13]. T3SS have been classified in 13 (I-XIII) major categories based on sequence divergence of T3SS core components [13].[13]According to this study, the *Vibrio* T3SS1 is part of category Ib, and the T3SS2 is the founding member of the T3SS phylogenetic category X [13].

Currently, the T3SS2 is considered the main virulence factor of *V. parahaemolyticus*. Studies in animal models of infection revealed that, while the T3SS1 plays a minor role during infection, the T3SS2 is essential for the enterotoxicity and the development of the clinical signs of the disease [14–17]. Therefore, the T3SS2 is considered the major virulence factor of pandemic *V. parahaemolyticus* [18]. The gene expression of the T3SS2 gene cluster is induced by bile salts [19, 20] through a regulatory network involving the VtrA, VtrC, and VtrB proteins [19, 21, 22] and the T3SS2 function requires host cell-surface fucosylation for efficient delivery of effector protein into infected cells [23]. Notably, the importance of the T3SS2 goes beyond increasing the pathogenic potential of pandemic strains, as the T3SS2 increases the environmental fitness of *V. parahaemolyticus* by promoting bacterial survival against predatory amoeba [24].

Phylogenetic analysis based on the amino acid sequence divergence of T3SS structural components has identified two distinct lineages of the T3SS2, known as phylotypes T3SS2α and T3SS2β [25, 26]. T3SS2α and T3SS2β gene clusters share a similar genetic organization and a subset of effector proteins (**Fig. 1**) [18]. A more recent study has proposed an additional category within T3SS2β, known as T3SS2γ [27]. The T3SS2γ is very similar to the T3SS2β gene cluster but has some gene content differences, such as the presence of a *tdh* gene inserted within the T3SS gene cluster [27] (**Fig. 1**).

**Fig. 1.**
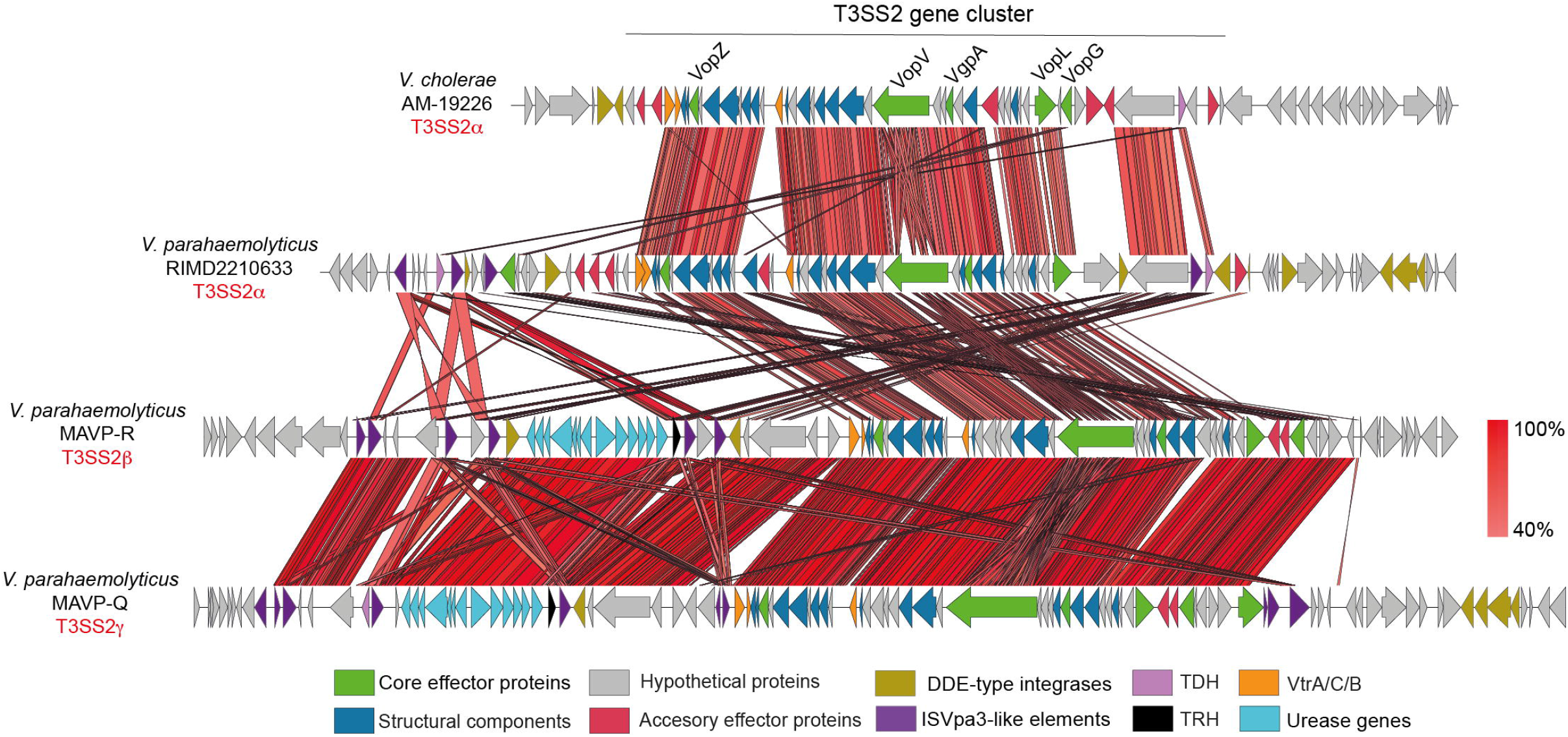
The T3SS2 is divided into two major phylotypes T3SS2α and T3SS2β, and a subcategory within T3SS2βknown as T3SS2γ. Schematic depiction of a comparison of the T3SS2 gene clusters in *V. cholerae* AM-19226, *V. parahaemolyticus* RIMD2210633, *V. parahaemolyticus* MAVP-R, and MAVP-Q. tBLASTx alignment was performed and visualized using Easyfig. Genes encoding different components are highlighted in color.

Most of our knowledge of the T3SS2 comes from studies of the T3SS2α of *V. parahaemolyticus* strain RIMD2210633 and the T3SS2α of *V. cholerae* strain AM-19226 (reviewed in [18, 28]). In this context, 15 different T3SS2 effector proteins have been identified, of which only 5 (VopZ, VopV, VopG, VopL and VgpA) are shared between *V. parahaemolyticus* RIMD2210633 and *V. cholerae* AM-19226 [29–43]. These 5 effectors are therefore called core effector proteins, while the remaining 10 are known as accessory effector proteins [18, 29, 39]. While useful, this classification has the limitation of comparing the distribution of effector of only 2 *Vibrio* strains of the same T3SS2 phylotype.

The T3SS2 gene cluster is not restricted to *V. parahaemolyticus* as similar T3SSs have been found in other species such as non-O1/non-O139 *V. cholerae, V. mimicus* and *V. anguillarum* [25, 44–47]. Furthermore, several reports have identified the presence of one or more T3SS2-related genes in strains of bacterial genera other than *Vibrio*, including *Aeromonas* [48], *Grimontia* [20], *Photorhabdus* [49, 50], *Providencia* [51] and *Shewanella* [29, 50]. Altogether, these reports suggest that the presence of T3SS2 gene clusters extends beyond the *Vibrionaceae* family and could have spread through HGT events. Nevertheless, these correspond to isolated observations performed in a limited number of strains and bacterial genomes. Therefore, the overall taxonomic and phylogenetic distribution of the T3SS2 and its effector proteins in bacterial genomes is currently unknown.

To address this issue, in this study, we performed a large-scale bioinformatic analysis to gain insight into the phylogenetic distribution of the T3SS2 gene clusters, including effector proteins. Our analysis identified 1130 bacterial genomes encoding putative T3SS2 gene clusters distributed over 47 different bacterial species from 8 bacterial genera. In addition, we classified these T3SS2s into 6 different subgroups in terms of the repertoire of effector proteins they possess. Our work provides evidence that the T3SS2 extends beyond the *Vibrionaceae* family and that acquisition of both the T3SS2 and different repertoires of effector proteins could have an impact on the pathogenic potential and environmental fitness of these bacteria.

### Impact Statement

One of the distinctive features of the O3:K6 pandemic clone of *Vibrio parahaemolyticus* is the presence of a phylogenetically distinct Type III Secretion System (T3SS) known as T3SS2. T3SS are complex nanomachines that allow the delivery of effector proteins directly into the cytosol of eukaryotic cells to manipulate different cellular functions. The *Vibrio* T3SS2 is the major virulence factor of the pandemic clone and a pivotal contributor to the pathogenesis and environmental fitness of the bacterium. Several studies have identified the T3SS2 gene cluster in other bacterial species beyond the *Vibrionaceae* family, suggesting that this T3SS can be mobilized through horizontal gene transfer events. In this work, we performed a phylogenetic distribution analysis of the T3SS2 gene cluster in publicly available bacterial genome sequences. We identified putative T3SS2 gene clusters in 1130 bacterial genomes from 47 different bacterial species from 8 genera and 5 bacterial families. Furthermore, we classified these T3SS2s into 6 different subgroups (I-VI) in terms of their repertoire of effector proteins. Our findings indicate that the T3SS2 is not restricted to the *Vibrionaceae* family and suggest that different effector protein repertories could have a differential impact on the pathogenic potential and environmental fitness of all the identified bacteria that have acquired this T3SS2.

## METHODS

### *In silico* identification and analyzes of T3SS2 loci and components

To identify putative T3SS2 loci, the amino acid sequence of the VPA1338 (SctN) and VPA1339 (SctC) proteins of *V. parahaemolyticus* RIMD2210633 were used as queries in BLASTp [52], to perform analyses using publicly available bacterial genome sequences of the National Center for Biotechnology Information (NCBI) database (February 2021). A cut-off of 70% identity and 80% sequence coverage threshold were used to select for positive matches. All genome sequence assemblies with positive hits were downloaded from the NCBI RefSeq or GenBank databases (See **Table S1** for accession numbers and metadata) and used to build local BLAST databases. To identify the remaining 35 T3SS2-related components in these 1130 genomes, the nucleotide and aminoacid sequences of each component (**Table S2**) were used as queries for either tBLASTx or BLASTp analyzes, respectively. A 50% identity and 80% sequence coverage threshold were used to select for positive matches. A threshold lower than the first screen was used to identify components which might have a higher sequence divergence. For comparative genomic analysis, nucleotide sequences were visualized and analyzed by the Artemis version 18.1 [53], the Artemis Comparison Tool (ACT) release 6 [54], the multiple aligner Mauve [55] and Easyfig v2.2.2 [56]. For the hierarchical clustering analysis, a presence/absence matrix of each T3SS2 component was constructed for each bacterial genome (**Table S1**) and uploaded as a *csv* file to the online server of the Versatile matrix visualization and analysis software MORPHEUS (one minus Pearson correlation, average linkage method) (https://software.broadinstitute.org/morpheus). The hierarchical clustering data analysis file from MORPHEUS is available as part of the online Supporting Data at the Zenodo data repository (https://doi.org/10.5281/zenodo.7016552). The presence and absence of each identified component were visualized as a colored matrix generated in GraphPad Prism version 9 (GraphPad Software, San Diego, California, USA).

### T3SS2 phylogenetic analyses

For T3SS phylogeny analysis, the 1130 concatenated SctN and SctC amino acid sequences were aligned with ClustalW using the Molecular Evolutionary Genetics Analysis (MEGA) software version 7.0 [57]. The phylogenetic tree was built from the alignments obtained from MEGA by performing a bootstrap test of phylogeny (1000 replications) using the Maximum-Likelihood (ML) method with a Jones-Taylor-Thornton (JTT) + CAT correction model using FastTree2 (Galaxy Version 2.1.10+galaxy1) [58] in the Galaxy platform (https://usegalaxy.org/) [59]. For the taxonomic phylogenetic analysis, a 16S RNA phylogenetic tree was constructed. The 16S rRNA DNA sequence of each bacterial species was obtained from the NCBI database and aligned with ClustalW using the Molecular Evolutionary Genetics Analysis (MEGA) software. The phylogenetic tree was built from the 16s rRNA DNA alignment by performing a bootstrap test of phylogeny (ultrafast bootsrap, 1000 replications) using the ML method using IQ-Tree [60](Galaxy Version 2.1.10+galaxy1). The phylogenetic trees were further visualized and edited to include important metadata with iTOL [61]. Files for the T3SS2 reconstructed phylogenetic tree (Newick file) is available as part of the online Supporting Data at the Zenodo data repository (https://doi.org/10.5281/zenodo.7016552). The T3SS2 phylogenetic tree can be interactively visualized in https://itol.embl.de/tree/19016190125374711626959067#.

### Identification of novel T3SS2 effector candidates

Each Open Reading Frame (ORF) within the predicted T3SS2 gene clusters of Subgroup VI (including 1 kb upstream and downstream of the first and last predicted component, respectively) was analyzed by means of the EffectiveDB [62], Bastion3 [63] and BEAN2.0 [64] pipelines to identify potential T3S signal sequences. ORFs with positive hits in at least one of these pipelines was further analyzed to predict putative functional domains with the InterProScan [65] and NCBI-CDD [66]. Prediction of other secretion signals such as for the Sec or TAT systems were analyzed with SignalP 6.0 [67]. Finally, each ORF was also analyzed by the structure-based homology tool HHpred [68]. Multiple sequence alignments were performed by MAFFT [69] and T-Coffee Expresso [70] and visualized by ESPript 3.0 [71].

## RESULTS AND DISCUSSION

### T3SS2 gene clusters extend beyond the *Vibrionaceae* family

We screened all publicly available bacterial genomes in the NCBI database to identify the full breadth of bacterial species harboring putative T3SS2 gene clusters. First, we performed BLASTp analysis in search for homologs of the VPA1338 (SctN) and VPA1339 (SctC) proteins of *V. parahaemolyticus* RIMD2210633. SctN and SctC were chosen as they are two of the most conserved T3SS structural proteins and are often used for phylogenetic clustering and classification of T3SSs [13, 72]. SctN is part of the ATPase complex and SctC is part of the export apparatus [18]. We used stringent cut-offs to avoid identifying homolog proteins from phylogenetically close T3SSs (≥70L% of identity and ≥80L% of coverage). Our analysis identified the presence of VPA1338 and VPA1339 homologs in 1130 bacterial genomes (**Fig. 2** and **Table S1**).

**Fig. 2.**
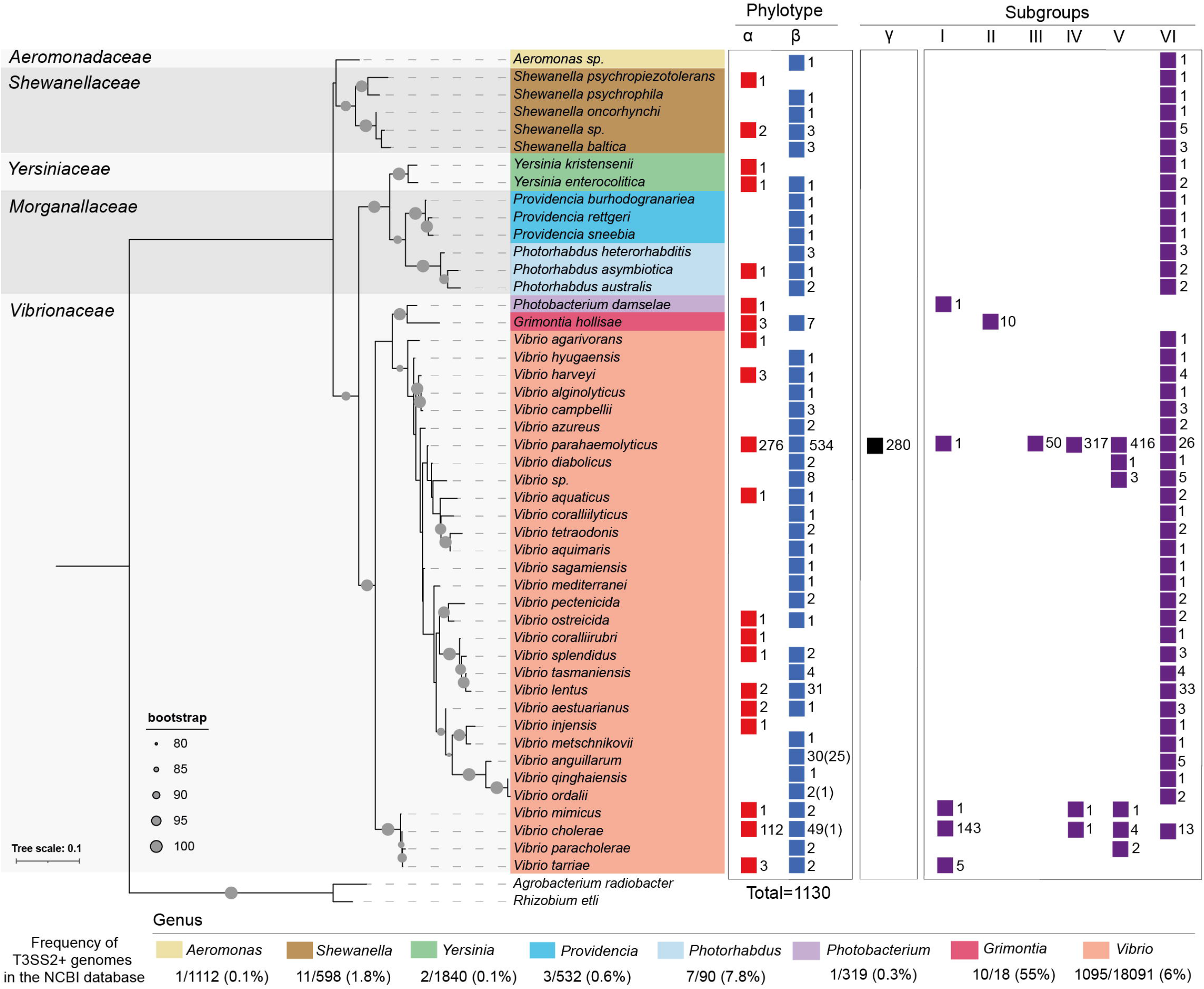
The T3SS2 extends beyond the *Vibrionaceae* family. Identification and taxonomic distribution of T3SS2 gene clusters in bacterial genomes. The presence of T3SS2 gene clusters from phylotypes T3SS2α, T3SS2β and different subgroups are shown for each taxonomic category (color-coded). The numbers correspond to the total numbers of bacterial genomes within each taxonomic category that harbor a putative T3SS2 gene cluster. In parenthesis the number of genomes lacking most T3SS2 structural components is shown. Maximum-likelihood phylogenetic tree was built from the alignments of 16S RNA DNA sequence from each representative bacterial specie with a bootstrap test of phylogeny (1000 replications) with IQ-Tree and visualized by iTOL. For each bacterial genus the percentage of T3SSS-positive bacterial genomes from the total NCBI database is shown.

To determine if these 1130 bacterial genomes could encode complete T3SS2 gene clusters, we further screened each genome for the presence of the 37 T3SS2-related components previously identified in *V. parahaemolyticus* RIMD2210633 (T3SS2α), *V. parahaemolyticus* MAVP-R (T3SS2β) and *V. cholerae* AM-19226 T3SS2α (**Table S1**).

These include ORFs encoding 16 structural proteins, 15 bacterial effector proteins, 3 regulatory proteins (VtrA, VtrB, and VtrC), and each of the 3 hemolysins linked to T3SS2 gene clusters (TDH_Vp_, TRH_Vc_ and TRH_Vp_). Most of the 1130 bacterial genomes (1106) encoded each of the T3SS structural genes suggesting that they most likely harbor complete T3SS2 gene clusters. The exception were 24 *V. anguilarum*, 1 *V. ordalii* and 1 *V. cholerae* genomes that only possess 8, 3 and 7 of the 16 T3SS2 structural components, respectively (**Table S1**). This was not unexpected as it has been previously reported that some *V. anguillarum* strains lack most of the T3SS2 structural components due to a large internal deletion of the T3SS2 gene cluster [73], suggesting that these strains do not encode functional T3SS2s.

Taxonomic distribution analysis identified putative T3SS2 gene clusters in 47 bacterial species from 5 distinct families (*Aeromonadaceae, Shewanellaceae, Yersiniaceae, Morganellaceae* and *Vibrionaceae*) and 8 bacterial genera (*Aeromonas, Shewanella, Yersinia, Providencia, Photorhabdus, Photobacterium, Grimontia* and *Vibrio*) (**Fig. 2**). In some genus, we identified multiple species harboring putative T3SS2 gene clusters while in other genus we could identify as little as one bacterial species harboring T3SS2 gene components (**Fig. 2**). In terms of bacterial species, the genus *Vibrio* contained the largest number of different species harboring putative T3SS2 gene clusters (31 species), followed by *Shewanella* (5 species), *Photorhabdus* and *Providencia* (each with 3 species), *Yersinia* (2 species) and the *Photobacterium, Grimontia*, and *Aeromonas* each with a single species. Finally, our analysis did not identify bacterial strains harboring more than one T3SS2 gene cluster within their genomes. Interestingly, several of these T3SS2 were identified in recently isolated and/or classified *Vibrio* and *Shewanella* species, including *V. paracholerae* [74], *V. tarriae* [75], *V. tetraodonis* [76], *S. oncorhynchi* [77] and *S. psychropiezotolerans* [78] (**Fig. 2** and **Table S1**).

Considering the total number of genome assemblies available in the NCBI database, putative T3SS2 gene clusters were more frequently identified in bacterial genomes from the *Grimontia* (55%), *Photorhabdus* (7.8%), *Vibrio* (6%) and *Shewanella* (1.8%) genus. With a lower frequency in genomes from the *Providencia* (0.6%), *Photobacterium* (0.3%), *Aeromonas* (0.1%) and *Yersinia* (0.1%) genus (**Fig. 2** and **Table S1**). While this suggests that the T3SS2 is overrepresented in some bacterial lineages, it could also reflect potential sequencing biases in terms of the diversity of bacterial genomics sequences deposited in the NCBI database (e.g. clinical versus environmental and different geographical origins).

Interestingly, most of the different bacterial species identified to harbor putative T3SS2 gene clusters can be found in aquatic habitats, suggesting that acquisition and transfer of the T3SS2 actively occur in this environment. Among the exceptions was the identification of T3SS2 gene clusters in genomes of strains of the *Photorhabdus* genus (**Fig. 2** and **Table S1**). Bacteria from the genus *Photorhabdus* comprise different bacterial species that are important insect pathogens, including some that can also cause human disease [79]. Wilkinson et al.[49], previously identified a T3SS2 gene cluster in the genome of *Photorhabdus asymbiotica* strain ATCC43949. Our analysis expanded this finding by identifying complete T3SS2 gene clusters in different strains of *P. australis, P. heterorhabitis* and *P. asymbiotica* species (**Fig. S1** and **Table S1**). Every T3SS2 gene cluster identified is inserted in the exact genomic location compared to the *P. laumondii* strain TT01 genome, as previously reported for *P. asymbiotica* strain ATCC43949 [49]. We also identified T3SS2 gene clusters in 1 strain of *Y. kristensenii* and 2 strains of *Y. enterocolitica* (**Fig. 2** and **Table S1**). Interestingly, while *Y. enterocolitica* can be found in aquatic environments, the 2 strains with putative T3SS2 gene clusters were isolated from wild ungulate carcasses and belong to the highly virulent biotype 1B [80].

Altogether, the taxonomic distribution of T3SS2 gene clusters in bacterial genomes indicates that this secretion system is not restricted to the *Vibrionaceae* family suggests that T3SS2 gene clusters can mobilize through HGT events among environmental bacteria. In support of this notion, it has been shown that under laboratory conditions naturally competent T3SS-negative *V. cholerae* strains can acquire and integrate the T332 gene cluster into their chromosomes suggesting that transfer of this gene cluster might occur in natural environments [81].

### The T3SS2β phylotype is most abundant and divergent than the phylotype T3SS2α

The T3SS2 is divided into two major phylotypes known as T3SS2α and T3SS2β [18], and an additional subcategory within T3SS2β known as T3SS2γ [27]. To determine the distribution of T3SS2α, T3SS2β and T3SS2γ, among the 1130 T3SS2s identified, we performed a phylogenetic analysis using concatenated SctN and SctC aminoacidic sequences. We also included the concatenated SctN and SctC sequences of the *Agrobacterium radiobacter* K84 and *Rhizobium etli* CIAT 652 T3SSs, which are representatives of T3SS category XI, the closest phylogenetic category to the *Vibrio* T3SS2 [13]. The phylogenetic clustering analysis showed that all the 1130 putative T3SS2s grouped closely together while the T3SS of *A. radiobacter* K84 and *R. etli* CIAT 652 cluster in a more distant branch (**Fig. 3**), supporting the notion that all the 1130 T3SS identified in this work correspond to bona fide T3SS2s.

**Fig. 3.**
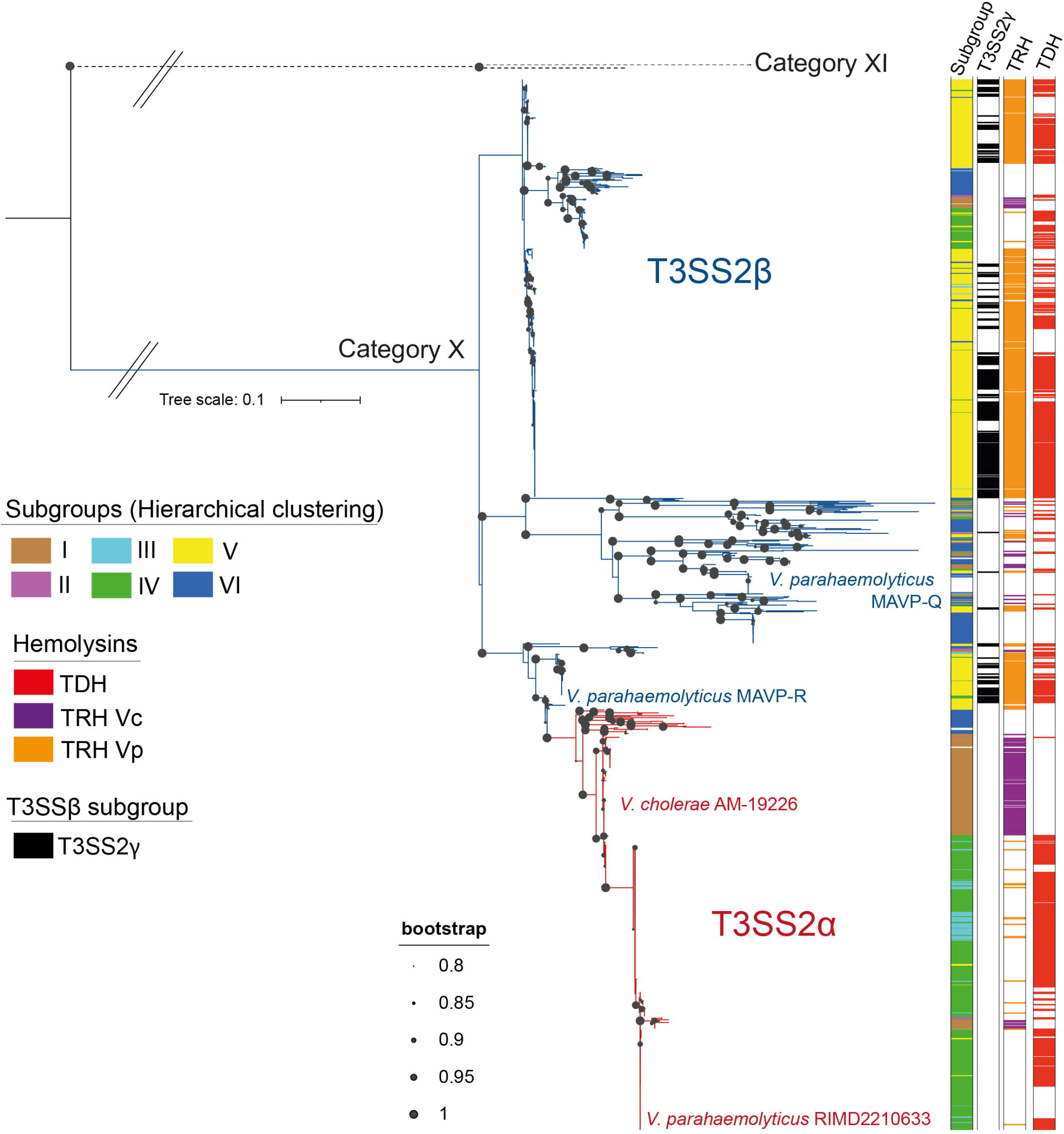
Phylogenetic analysis of the identified T3SS2s. Maximum-likelihood phylogenetic tree built from the alignments of concatenated SctN and SctC amino acid sequences with a bootstrap test of phylogeny (1000 replications) with a Jones-Taylor-Thornton (JTT) correction model and visualized by iTOL. The presence of hemolysis and subgroups are shown as colored strips. T3SS2α and T3SS2β are shown in red and blue branches, respectively. The bacterial strains of representative T3SS2α and T3SS2β Gene clusters are also highlighted in red and blue color, respectively.

To classify these T3SS2 in terms of phylotypes (α or β), we used the previously classified T3SS2α of *V. cholerae* AM-19226 and *V. parahaemolyticus* RIMD2210633 and the T3SS2β of *V. parahaemolyticus* MAVP-Q and MAVP-R strains to identify the major two branches that separate these two phylotypes (**Fig. 3**). Our analysis also revealed that 36,7% (n= 415) of the identified T3SS2s belonged to phylotype T3SS2α and 63,3% (n= 715) to phylotype T3SS2β. The phylogenetic analysis also revealed that the greater sequence divergence was observed within the T3SS2β phylotype (**Fig. 3**).

As mentioned above, recent reports have created a subcategory within T3SS2β of *V. parahaemolyticus* known as T3SS2γ (based mainly on the presence of both the *tdh* and *trh* genes) [27]. To determine the percentage of T3SS2γ gene clusters within the T3SS2β family, we performed a gene content analysis based on the definition of T3SS2γ gene clusters proposed by Xu et al., [13]. The analysis identified that 40,5% (n= 280) of T3SS2β gene clusters could be classified as T3SS2γ. Notably, the presence of T3SS2γ did not cluster to any major clade of the T3SS2 phylogenetic tree (**Fig. 3**), supporting the notion that T3SS2γ is not a third T3SS2 phylotype but rather a subcategory within T3SS2β.

Taxonomic distribution analysis identified the presence of T3SS2α and/or T3SS2β in strains of each of the 5 bacterial families (**Fig. 2 and Fig. 3**), with no clear overrepresentation of any phylotype within a particular bacterial family. Nevertheless, T3SS2β was present in a more significant number of different bacterial species (41 out of 47) in comparison to T3SS2α (20 out of 47 species) (**Fig. 2**). Finally, the T3SS2γ subcategory was identified solely in *V. parahaemolyticus* strains which agree with previous reports [27]. Since many *V. parahaemolyticus* strains which harbor T3SS2 gene clusters are related to the RIMD2210633 pandemic clone, we analyzed the distribution of T3SS2 phylotypes among different *V. parahaemolyticus* Sequence Types (STs). Our analysis showed that T3SS2α was overrepresented in *V. parahaemolyticus* strains from ST3 (44%), ST120 (11,5%), ST189 (7%) and ST332 (16,4%) while the T3SS2β was overrepresented in strains from ST3 (11,5%), ST36 (25,2%) and ST631 (7,5%) (**Fig. S2**).

### T3SS2 gene clusters share a high degree of synteny

To gain insight into the genomic context of the identified T3SS2 gene clusters, we performed a comparative genomic analysis between the T3SS2 gene cluster of *V. parahaemolyticus* RIMD2210633 and representative strains of 74 non-*Vibrio* bacterial species (**Table S1**) with complete or almost complete bacterial genomes (large enough contigs to allow a comparative analysis). Notably, there was a high gene synteny between the T3SS2 gene cluster of *V. parahaemolyticus* RIMD2210633 and the gene clusters of bacterial species of different genera (e.g., *Providencia, Photobacterium, Grimontia* and *Photorhabdus*) (**Fig. 4A, Fig. 4B, Fig. 5A** and **Fig. S1**). In each of the T3SS2 gene clusters identified in *Grimontia hollisae* strains, a large gene encoding a putative adhesin (SiiE-like adhesin) is encoded in the middle of the T3SS2 gene cluster (**Fig. 5A**). A high gene synteny was also observed between the T3SS2 gene clusters with a lower sequence identity, such as the gene clusters of *Aeromonas* sp. DNP9 and *Yersinia enterocolitica* Y1Wildboar1B (**Fig. 4C**).

**Fig. 4.**
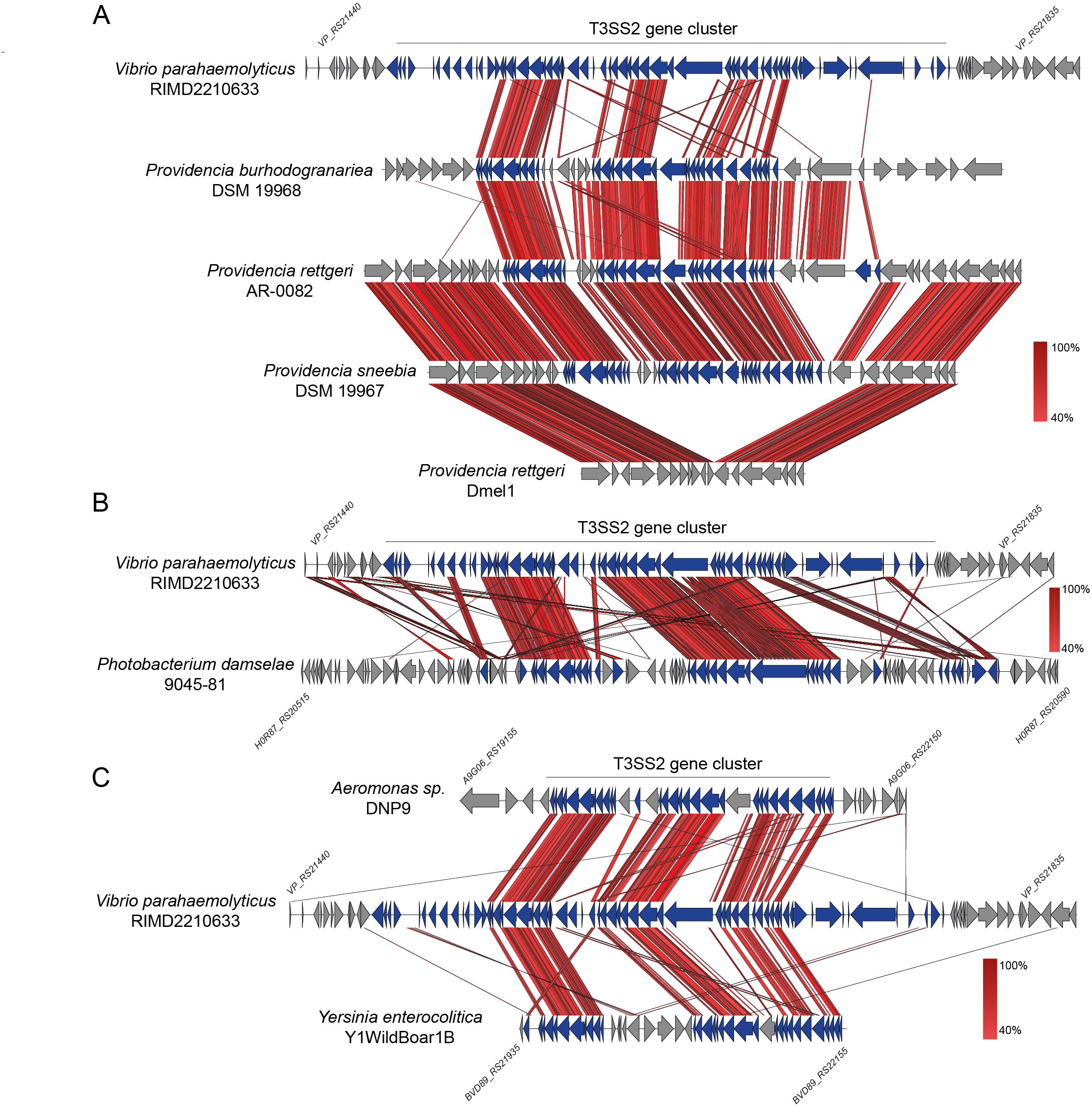
T3SS2 gene clusters share a high degree of synteny. Schematic depiction of a comparison of the T3SS2 gene clusters in *V. parahaemolyticus* RIMD2210633 in comparison to the gene clusters of strains of *Providencia* (A), *Photobacterium* (B) and *Aeromonas*, and *Yersinia enterocolitica* strains (C). tBLASTx alignments were performed and visualized using Easyfig. Genes encoding T3SS-related genes are highlighted in blue.

**Fig. 5.**
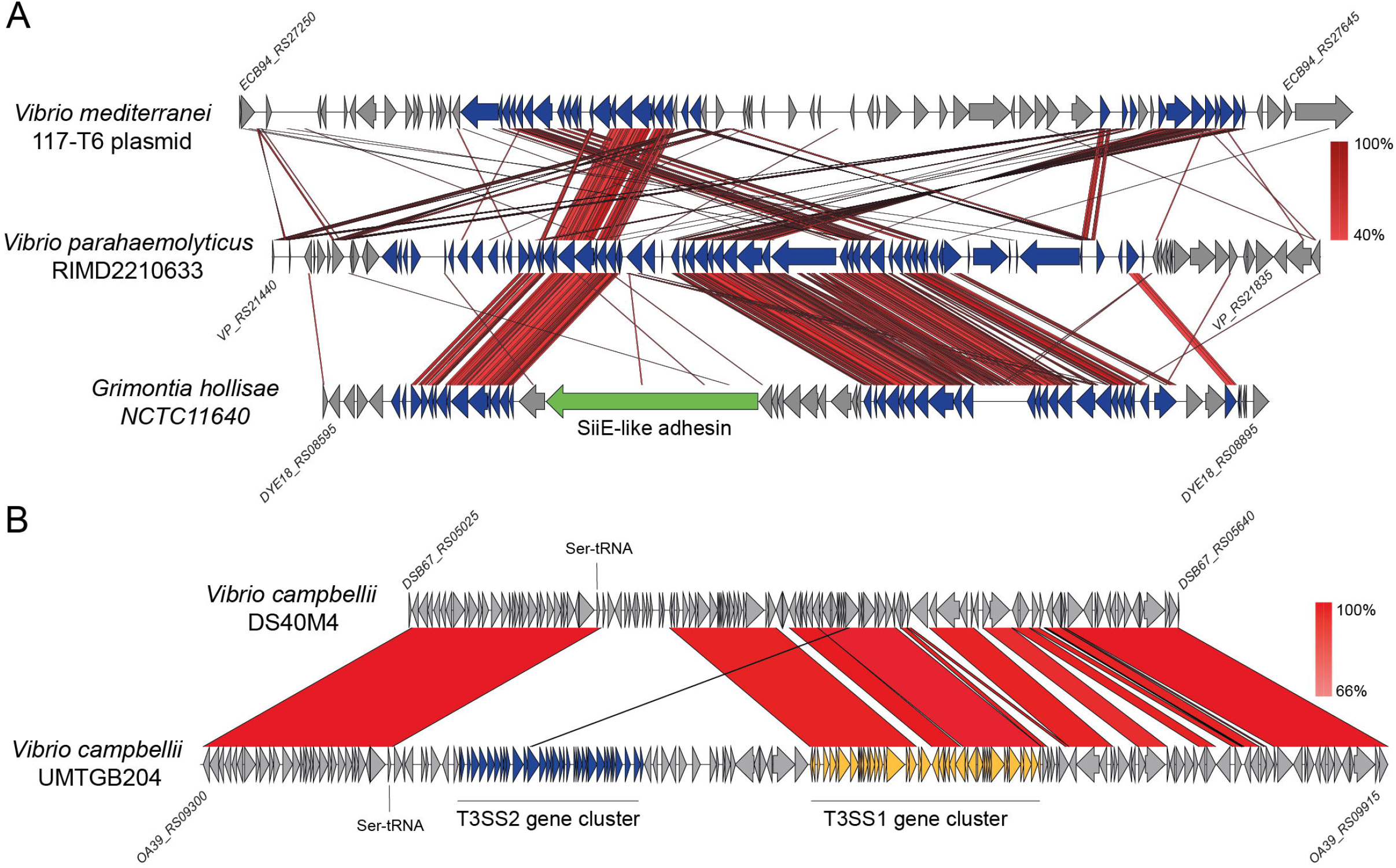
Distinctive genomic locations of T3SS2 gene clusters. (A) Comparison of the T3SS2 gene cluster in *V. parahaemolyticus* RIMD2210633 with the T3SS2 gene cluster encoded in *V. mediterranei* 116-T6 plasmid and the T3SS2 gene cluster of *G. hollisae* NCTC11640. tBLASTx alignmens was performed and visualized using Easyfig. (B) Schematic depiction of a comparison of the T3SS1 and T3SS2 gene clusters inserted in the vicinity of the serine tRNA in *Vibrio campbellii* strain UMTGB204. BLASTn alignments was performed and visualized using Easyfig. Genes encoding T3SS-related genes are highlighted in blue.

For *Vibrio* species, we also analyzed the genome location of these T3SS2 gene clusters. In addition to the described chromosomal location of T3SS2 within VPAI-7 of *V. parahaemolyticus* RIMD2210633 [4], we identified a T3SS2 gene cluster within the plasmid of a recently sequenced *V. meditterranei* strain 117-T6 (CP033579) [82] (**Fig. 5A**). Another interesting observation was the presence of a T3SS2 gene cluster inserted right next to a T3SS1 gene cluster in *V. campbellii* strain UMTGB204. In this strain both T3SSs are inserted within the vicinity of the same serine tRNA (**Fig. 5B**), in which the T3SS1 of *V. parahaemolyticus* RIMD2210633 is located.

### Hierarchical clustering analysis identifies 6 subgroups with different repertoires of effector proteins

While phylogenetic analysis of T3SSs using core structural components provides insight into the evolution of T3SSs, the acquisition, transfer, or loss of accessory and effector encoding genes provides an additional layer of complexity to the evolution of these systems [11, 13, 72]. To gain further insight into this diversity, we performed a hierarchical clustering analysis of 37 T3SS2-related genes in the 1130 bacterial genomes (**Fig. 6**). For the analysis of the effector proteins, we included each T3SS2 effector described for either *V. parahaemolyticus* RIMD2210633 or *V. cholerae* AM-19226 [18]. The hierarchical clustering analysis allowed us to identify 6 major subgroups (I to VI) with similar gene content profiles (**Fig. 6A** and **Table S1**).

**Fig. 6.**
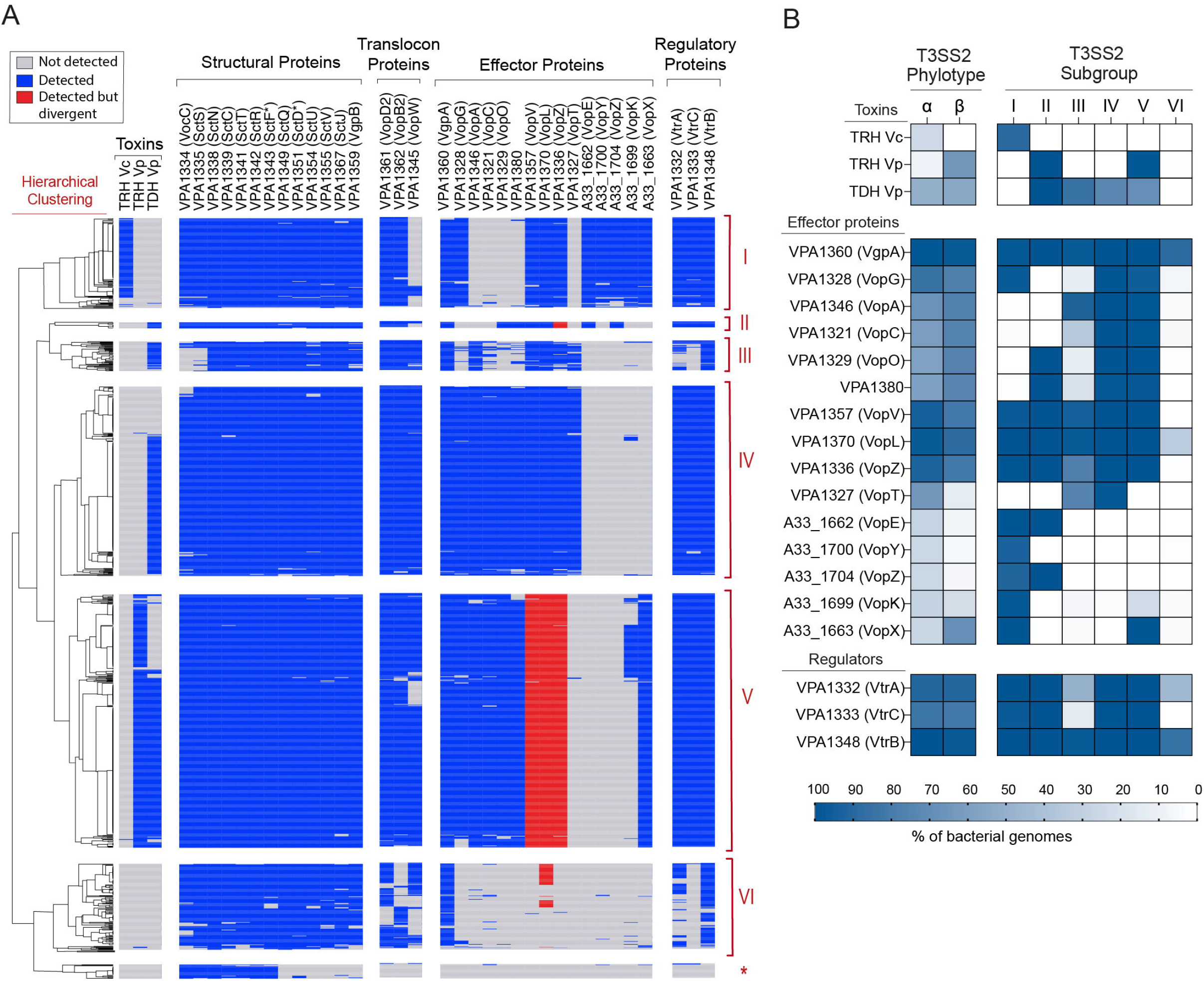
Hierarchical clustering analysis of T3SS2-related genes identifies 6 distinct subgroups. (A) Clustering analysis was performed using Versatile matrix visualization and analysis software MORPHEUS (one minus Pearson correlation, average linkage method). Boxes representing genes not detected, detected, or detected but divergent (less than 70% amino acid sequence identity with at least 80% sequence coverage) are colored in grey, blue or red, respectively. An asterisk highlights the group of *V. anguillarum* genomes that only harbored a limited number of T3SS structural components. (B) Comparison of the frequency of detection of T3SS2 effector proteins, toxins and regulators between each T3SS2 phylotype and subgroup. Frequencies are color-coded and expressed as percentages.

Most structural components were highly conserved in all subgroups, except for the hydrophilic translocator VopW of subgroups I and VI, the structural component SctS and the VocC chaperone of subgroup II, and the VopB2 and VopD2 translocon proteins of subgroup VI (**Fig. 6A**). While it is possible that these gene clusters do not encode such components, the inability to identify these components could be due to the lower sequencing coverage (<68X) of most of these bacterial genomes. The *vtrA, vtrB*, and *vtrC* genes, which encode the regulatory complex that induces the expression of T3SS2 gene clusters in response to bile [19, 22], were also highly conserved, except for subgroups III and VI, where many bacterial strains harbored divergent homologs of VtrA and VtrC (**Fig. 6A**).

Since most structural genes were highly conserved across all genomes, the hierarchical clustering analysis mainly classified each T3SS2 based on the repertoire of effector proteins identified. As shown in **Fig. 6B**, each T3SS2 subgroup was enriched in a distinct set of effector proteins and associated hemolysins. While most subgroups were enriched for a combination of 10-12 different T3SS2 effectors (subgroups I to V), subgroup VI was vastly underrepresented in terms of the presence of known effector proteins. This subgroup was primarily enriched in the VgpA effector protein, lacking most other known T3SS2 effector proteins. The presence of VgpA homologs in each subgroup supports this effector’s critical role in T3SS2 function [39, 43]. As expected, the current phylotype classification (T3SS2α versus T3SS2β) was a poor predictor of the presence/absence of individual effector proteins (**Fig. 6A** and **B**), highlighting the utility of a gene-content classification scheme.

Interestingly, most T3SS2 subgroups were enriched for genomes of particular bacterial species (**Fig. 2** and **Table S1**): i) Subgroup I included strains from 5 different bacterial species (*P. damselae, V. cholerae, V. mimicus, V. parahaemolyticus*, and *V. tarriae*.); ii) Subgroup II included strains only from *G. hollisae* species; iii) Subgroup III included *V. parahaemolyticus* strains exclusively; iv) Subgroup IV included strains from 3 species (*V. cholerae, V. mimicus* and *V. parahaemolyticus*); v) Subgroup V included strains from 6 species (*V. parahaemolyticus, V. cholerae, V. diabolicus, V. mimicus, V. paracholerae* and *Vibrio sp*.); and subgroup VI harbored the most diverse set of bacterial species with a total of 42 bacterial species (**Fig. 2**). Subgroup VI was also the only subgroup with bacterial strains beyond the *Vibrionaceae* family. Finally, while no T3SS2 subgroup was exclusive to a particular T3SS2 phylotype some subgroups were indeed overrepresented by different phylotypes. Subgroups I, III and IV were overrepresented in strains from the T3SS2α phylotype, while subgroups II, V and VI were overrepresented in strains from the T3SS2β phylotype (**Fig. 3** and **Table S1**). Finally, every *V. parahaemolyticus* strain of the T3SS2γ subcategory clustered within subgroup V (**Table S1**).

Altogether, the hierarchical clustering analysis suggests that T3SS2s have acquired a diverse set of effector proteins which could ultimately impact on the contribution of each T3SS2 to the pathogenic potential and environmental fitness of different bacterial strains.

### T3SS2 subgroup VI encodes 10 novel effector candidates

As mentioned above, the T3SS2 gene clusters of subgroup VI stand out as they are the most taxonomically diverse (**Fig. 2**) and lack most of the known T3SS2 effector proteins (**Fig. 6**). The high degree of synteny of these gene clusters to the T3SS2 gene cluster of *V. parahaemolyticus* RIMD2210633 (**Fig. 4 and Fig. S1**) and their distribution in the T3SS2 phylogenetic tree (**Fig. 3**) suggests that these are not part of a phylogenetically distinct T3SS2, but rather have a different set of effector proteins. To test this hypothesis, we analyzed the vicinity of the T3SS2 gene cluster from 1 strain of each species of subgroup VI (**Table S1**) and performed bioinformatic analysis to identify putative T3SS2 effector candidates. We analyzed each ORF of these T3SS2 clusters based on 4 approaches, including: i) identification of putative T3SS secretion signals by T3SS effector prediction pipelines (BEAN2.0, Bastion3 and EffectiveDB); ii) bioinformatic analysis to identify conserved functional domains (InterProScan, NCBI-CDD); iii) sequence identity to known T3SS effector proteins of other bacterial species (BLASTp, amino acid sequence identity >40%); and iv) functional predictions via remote homology searches using the HHpred pipeline.

Our analysis identified a set of 10 novel effector candidates that harbored putative T3SS secretion signal sequences and putative functional domains present in known T3SS effector proteins. These included putative ADP-ribosyltransferases, tyrosine phosphatases, glycosyltransferases and Rho-activating proteins (**Table 1, Fig. 7** and **Fig. S3**). In addition, 5 of these ORFs encode homologs of known T3SS effector proteins or toxins such as the effector proteins OspC1, AoPH and VopC of *Shigella flexneri, Aeromona hydrophila* and *V. parahaemolyticus* respectively (**Table 1, Fig. 7** and **Table S2**). The 10 effector candidates identified were present in diverse bacterial species of the T3SS2 subgroup VI, except for the putative tyrosine phosphatases (DYE18_RS08865) that also was identified in *Grimontia hollisae* strains of T3SS2 subgroup II (**Fig. 7** and **Table S2**).

**Table 1.** Prediction of T3SS effector candidates within the T3SS2 subgroup VI. Representative ORFs for each prediction are shown with the results of different prediction software (1: BEAN2.0; 2: BastionX; 3: EffectiveT3; 4: EffectiveCCBD; 5: EffectiveELD), the identification of putative functional domains (InterproScan), and the amino acid sequence identity to known T3SS effector proteins from other organisms. ND, not determined.

| Effector Candidate | Length (aa) | T3SS effector prediction | Functional domain Prediction | Similarity to known T3SS effectors or toxins |
| --- | --- | --- | --- | --- |
| OA39_RS25565 | 317 | 1,2,3,4 | Tyrosine phosphatase (IPR029021) | ND |
| ECB94_RS27680 | 409 | 1,2,3,4 | CNF1 Rho-activating domain (IPR008430) | 27,7% identity to VopC (blastp, 3e-48 Evalue) |
| ECB94_RS27290 | 478 | 1,2,3,4 | Shigella flexneri OspC (IPR010366) | 49,9% identity to OspC1 (blastp, 5e-153 Evalue) |
| Ppb6_RS09600 | 472 | 1,3,4 | ADP-ribosylation (SSF56399) | ND |
| CGJ34_RS08520 | 320 | 1,3,4 | Glycosyltransferase (IPR029044) | Similarity to Lgt1 toxin (HHpred, 2.4e-16 Evalue) |
| A9G06_RS19230 | 425 | 1,2 | ADP-ribosylation (SSF56399) | ND |
| CGJ46_RS22230 | 403 | 1,4 | ADP-ribosylation (SSF56399) | ND |
| BCT80_RS20435 | 400 | 1 | GDSL lipase (IPR001087) | 22,4% identity to VPA0226 (blastp, 3e-18 Evalue) |
| DYE18_RS08865 | 401 | 1,5 | Tyrosine phosphatase SptP/YopH (IPR003546) | 61,2% identity to AopH (blastp, 1e-119 Evalue) |
| HUW04_RS16370 | 283 | 1,5 | Rho GTPase-activating domain (IPR000198) | ND |

**Fig. 7.**
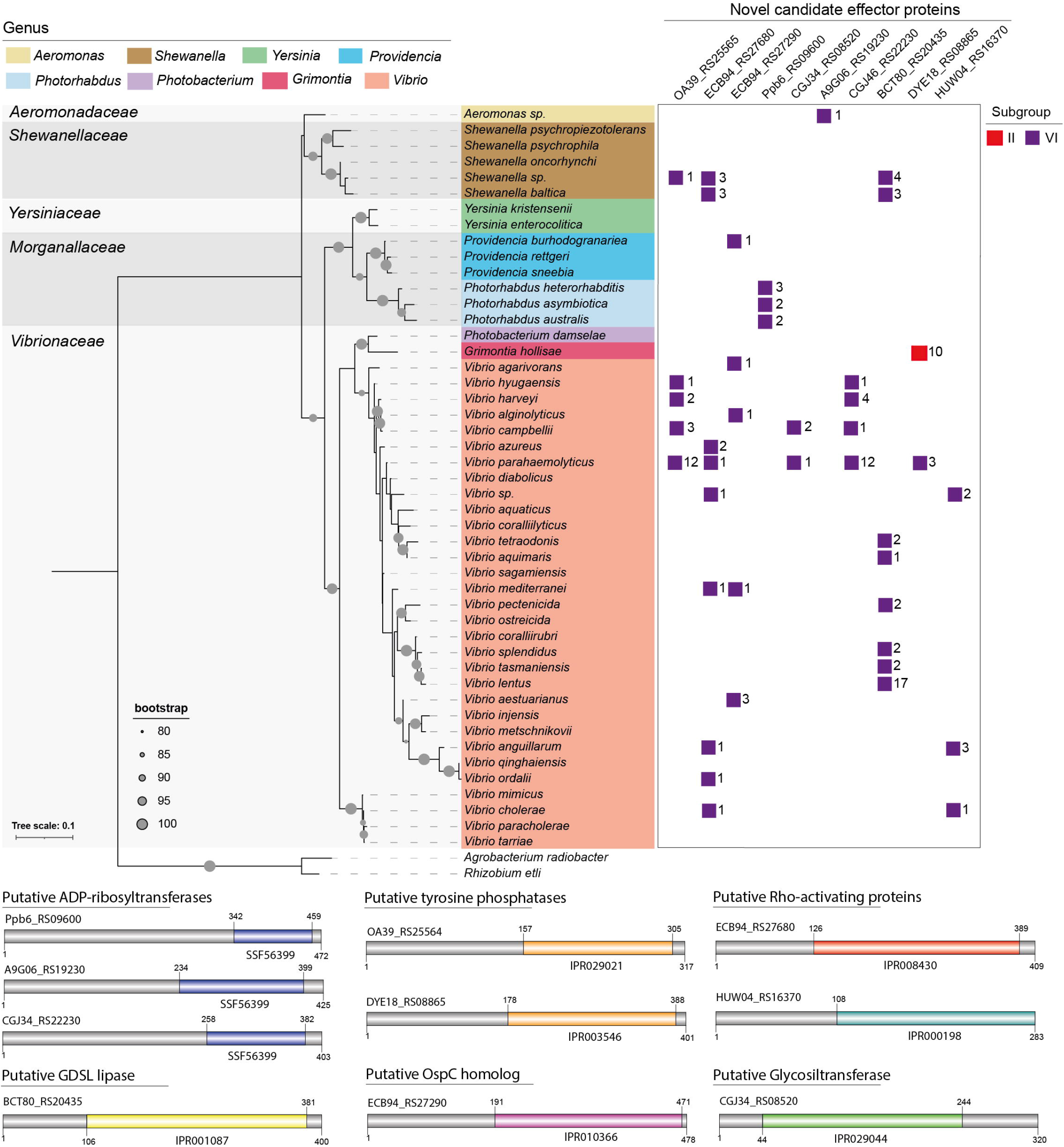
T3SS2 Subgroup VI encodes 10 novel effector candidates. Taxonomic distribution of each predicted effector protein identified in bacterial genomes. The numbers correspond to the total numbers of bacterial genomes within each taxonomic category that harbor a novel effector candidates. Schematic representation of novel candidate highlighting the presence of predicted functional domains in different colors is shown below.

Interestingly, one candidate (BCT80_RS20435) harbored a predicted GDSL lipase domain (IPR001087) and a 22,4% amino acid sequence identity to the VPA0226 protein of *parahaemolyticus* (**Fig. S4**). VPA0226 is a secreted lipase required for *V. parahaemolyticus* RIMD2210633 to egress from the infected cell. Notably, in *V. parahaemolyticus* VPA0226 is not secreted by the T3SS2 but rather by the T2SS [83]. Unlike VPA0226, analysis of BCT80_RS20435 showed no detectable T2SS or TAT secretion signal (**Fig. S4**).

Overall, our analysis suggests that subgroup VI encodes a subset of novel T3SS2 effector proteins. Further work is needed to confirm that each candidate corresponds to T3SS2 effector proteins.

### Revisiting the current classification of core and accessory T3SS2 effector proteins

Recently, Matsuda et al. [18] proposed a classification of T3SS2 effector proteins based on the presence of these effectors in *V. parahaemolyticus* and *V. cholerae*.

Effectors present in both *V. cholerae* and *V. parahaemolyticus* are known as core effectors, and those present in either *V. cholerae* or *V. parahaemolyticus* have been called accessory effector proteins. While useful, this classification relies on analyzing only two strains of two bacterial species.

To revisit this classification, we analyzed the presence/absence of each T3SS2 effector protein in the 47 bacterial species in which we have identified putative T3SS2 gene clusters. As shown in **Fig. S5**, only VgpA was present in at least one strain of most bacterial species (except for *Y. enterocolitica* and *Y. kristensenii* strains). The high prevalence of this effector protein could be explained by its additional structural role [39]. Another effector, VopT, was identified exclusively in *V. parahaemolyticus* strains. The remaining 13 T3SS2 effector proteins were present at various degrees with no overrepresentation in any bacterial species. Altogether, our data show a wide distribution of T3SS2 effector proteins in different bacterial species and that there is no clear correlation between the number of bacterial species that harbor a particular effector protein, which is the basis of the current classification scheme. We argue that a better alternative is to classify T3SS2s based on the repertoire of their effector proteins in the context of all identified T3SS2 loci in bacterial genomes (subgroups I-VI).

Nevertheless, such a classification scheme is not without limitations. First, we believe that for such a classification scheme to be useful a larger dataset of T3SS2-positive bacterial genomes beyond the *Vibrionaceae* family are needed, to fully grasp the diversity of this system. Second, a large fraction of these genomes would need to be of high quality to avoid sequence coverage bias. And finally, an inherent limitation is that this classification scheme could turn out to be dynamic, as it may change over time if novel effector encoding genes are acquired through HGT or if some are lost due to genomic rearrangements.

It has been shown that genes encoding bacterial effector proteins can be acquired through HGT and that can also further evolve through terminal reassortment and pseudogenization events [11, 12, 84–86]. This supports the notion that different repertoires of effector proteins will be selected by different bacterial species in terms of their contribution to their adaptation to different ecological niches. In this context, it could be possible that different T3SS2 positive bacterial strains use different effector protein repertoires (subgroups I-VI) to differentially interact with environmental eukaryotic hosts which ultimately would impact on their pathogenic potential.

## CONCLUSION

The acquisition and transfer of T3SS gene clusters can potentially increase the virulence and the environmental fitness of the recipient bacteria. In this context, it has been proposed that the acquisition of the T3SS2 gene cluster by *V. parahaemolyticus* was a crucial factor in the global spread and increased pathogenic potential of the O3:K6 pandemic clone and its derivatives. In this study, we performed a genome-wide analysis to determine the phylogenetic distribution of the *Vibrio* T3SS2 gene cluster in public genome databases. Our analysis revealed that T3SS2 gene clusters extend beyond the *Vibrionaceae* family, share a high degree of synteny and that it is possible to categorize T3SS2 in 6 different subgroups (I-VI) in terms of their repertoire of effector proteins.

Notably, it is plausible to think that acquisition of the T3SS2 gene cluster by HGT events in each of the bacteria identified in this study could increase their human pathogenic potential and environmental fitness, just as it has been proposed for the *V. parahaemolyticus* pandemic clone. Finally, since the repertoire of T3SS effector proteins delivered to the target cells ultimately defines the cellular outcome of an infection, we believe that future studies should focus on testing the contribution of each repertoire of T3SS2 effector proteins to the pathogenic potential and environmental fitness of these bacterial species.

## AUTHOR CONTRIBUTIONS

Sebastian Jerez: Investigation, Visualization, review and editing final manuscript.

Nicolas Plaza: Investigation, Visualization, review and editing final manuscript.

Italo M Urrutia: Investigation, Visualization, review and editing final manuscript.

Veronica Bravo: Investigation, Visualization, review and editing final manuscript.

Carlos J Blondel: Conceptualization, Methodology, Formal analysis, Investigation, Visualization, Resources, Data curation, Writing-Original Draft and review and editing., Funding acquisition, Supervision and Project administration.

## ACKNOWLEDGMENTS

We thank members of the Blondel Laboratory for helpful discussions on all aspects of this project and for their comments on the manuscript.

## FIGURE LEGENDS

**Table S1**. Table with information regarding the presence/absence and hierarchical clustering analysis of T3SS2 components in the bacterial genome sequences (including accession numbers).

**Table S2**. Table with information regarding the novel putative effector candidates.

**Fig. S1**. Gene synteny between the T3SS2 gene clusters of *V. parahaemolyticus* RIMD2210633 and the identified T3SS2 gene clusters of *Photorhabdus* species. tBLASTx alignments were performed and visualized using Easyfig. Genes encoding T3SS-related genes are highlighted in blue.

**Fig. S2**. Distribution of *V. parahaemolyticus* Sequence Types (STs) in terms of the presence of T3SS2α or T3SS2β gene clusters.

**Fig. S3**. Genetic context of novel effector candidates in representative T3SS2 gene clusters. The T3SS2 gene clusters are shown in blue and each novel effector candidate is highlighted in red.

**Fig. S4**. Multiple sequence alignment of the amino acid sequence of the predicted effector BCT80_RS20435 and the secreted lipase VPA0226 and the *Salmonella* T3SS effector protein SseJ. The alignment was performed using T-Coffee Expresso and visualized by ESPript 3.0. Amino acids within a red background correspond to positions with 100% identity; amino acids with a yellow background correspond to positions with >70% identity SignalP analysis shows that BCT80_RS20435, unlike VPA0226, does not harbor Sec or TAT secretion signals.

**Fig. S5**. Distribution of T3SS2 effector proteins among bacterial species. The distribution matrix shows the detection of each effector protein in at least one bacterial strain in red. The phylogenetic tree is the same tree as Fig 2.

