## Supplementary figures and images for "The *Vibrio* Type III Secretion System 2 is not restricted to the *Vibrionaceae* and encodes differentially distributed repertoires of effector proteins"

### Fig. S1

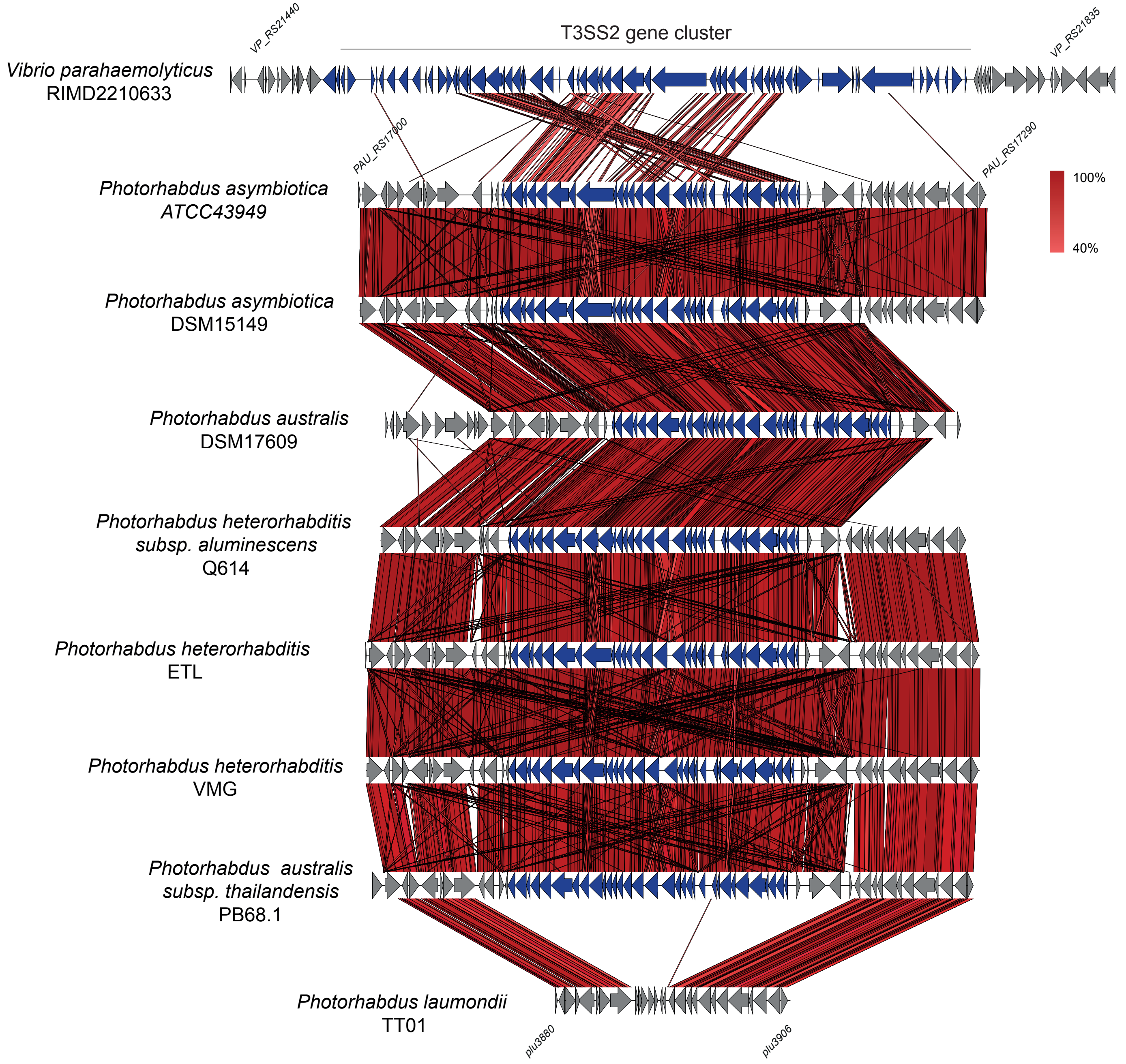

### Fig. S2

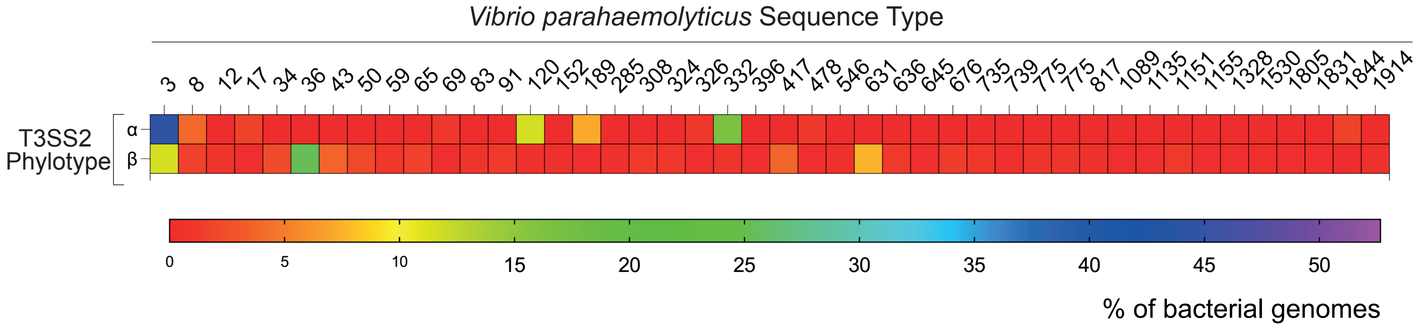

### Fig. S3

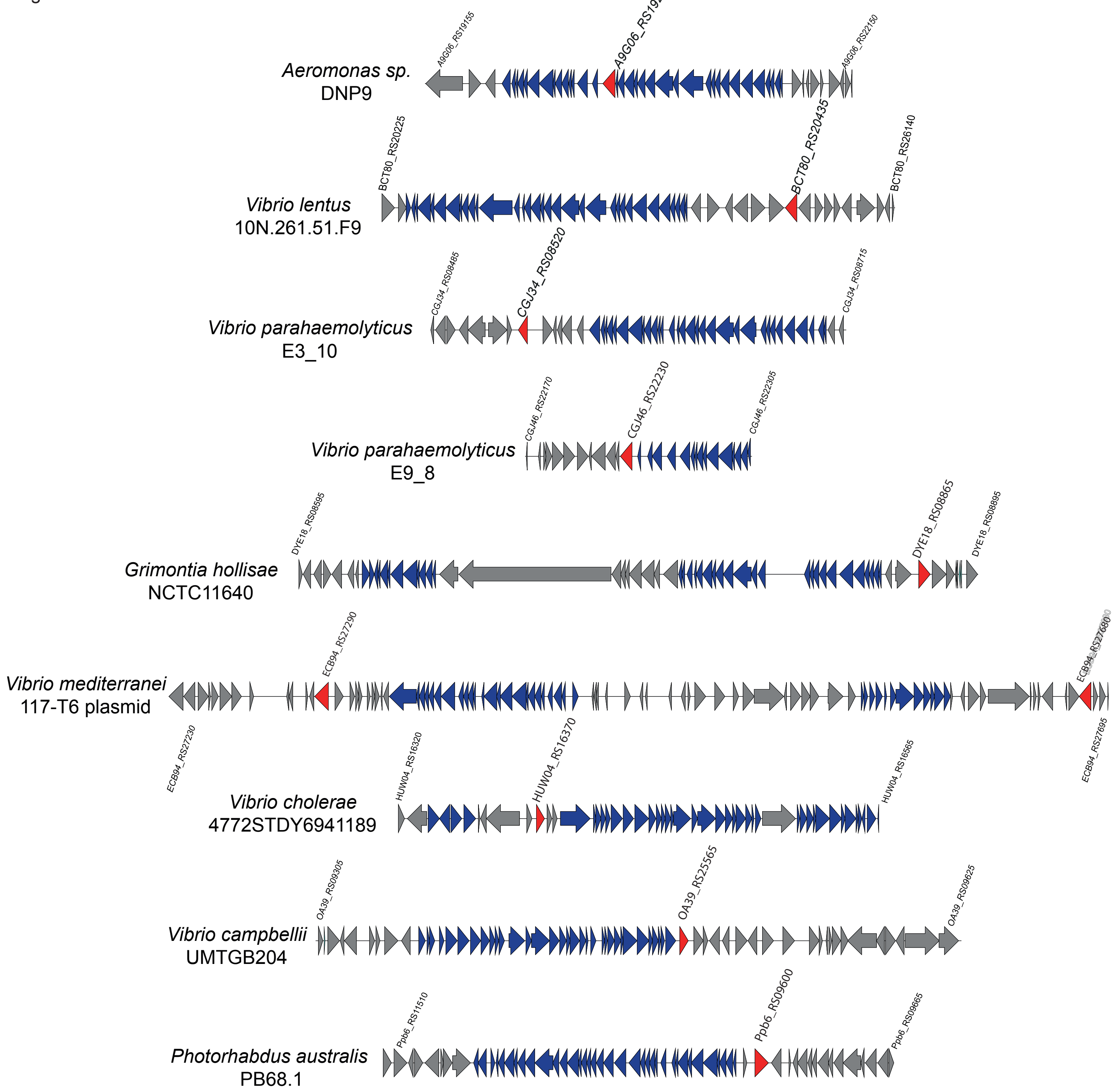

### Fig. S5

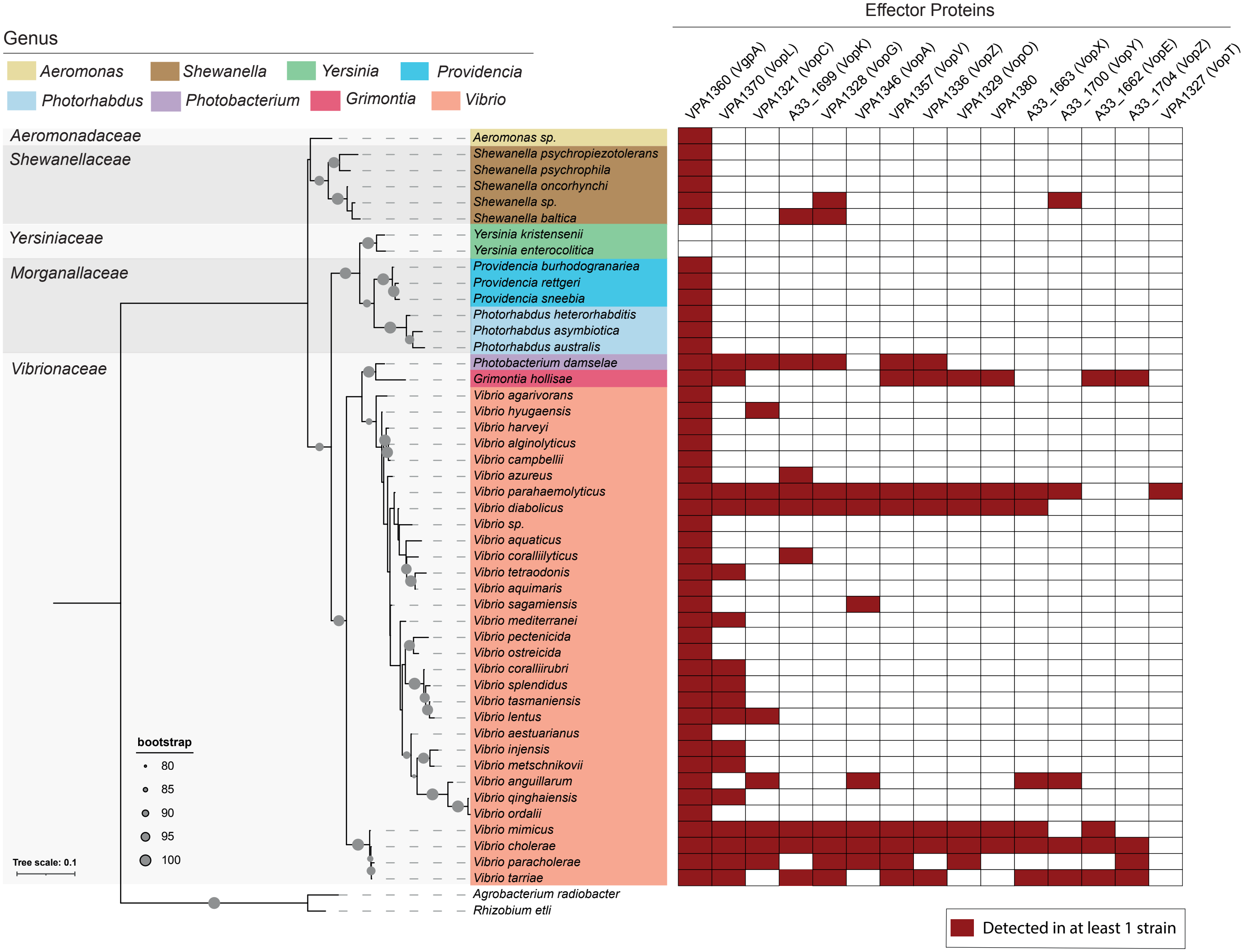
