## Supplementary material for "The *Vibrio* Type III Secretion System 2 is not restricted to the *Vibrionaceae* and encodes differentially distributed repertoires of effector proteins": Fig. S4

```
                1         10        20        30        40
BCT80_RS20435   MKS..LILGRQL...VS.IEYNKKLSERKVKEEKIRVAKVSQLSSYDKLKTPVLV...............
VPA0226         MMKKTITLLTALLPLASAVAEEPTLSPEMVSAS.E.VISTQENQTYTYVRCWYRTSYSKDDPATDWEWAK
sseJ_           MPL...SVGQGY........FTSSISSEKFNAIKE.SARLPELSLWEKIKAYFFTTHHAEALECIFNLYH
consensus>70    M........................S...............#............................

                50        60        70        80        90
BCT80_RS20435   ...................LNEKVEH..VIDGTELGFDS...TDGAEHKDLLFELAIRNKE....VTNAQ
VPA0226         NEDGSYFTIDGYWWSSVSFKNMFYTNTSQNVIRQRCEATLDLAN..ENADITFFAADNRFSYNHTIWSND
sseJ_           HQELNLTPVQV........RGAYIKL..RALASQGCKEQFI.IESQEHADKLIIKDDNGEN....ILSIE
consensus>70    .................................#.........#..E..D...............!...#

                100       110       120       130       140
BCT80_RS20435   L.................TSHSSQALDHIKRLVVIGDSLSDSEGRMKSKTLGVMLSSRQYYKGRFTNGFV
VPA0226         A.................AMQPDQ....INKVVALGDSLSDT.GNIFNASQWRFPNPNSWFLGHFSNGFV
sseJ_           VECHPEAFGLAKEINKSHPKPKNISLGDITRLVFFGDSLSDSLGRMFEKTHHILPSYGQYFGGRFTNGFT
consensus>70    ............................I...V..GDSLSD..G................%.G.F.NGF.

                150       160       170       180       190       200       210
BCT80_RS20435   WADFISSGAYLKKHMKSEEARDNFKLLNYAEGGAVT.AKYSKLNPTFWFISNMK...RQIHKHEKKEGFQ
VPA0226         WTEYIAKAKNL.............PLYNWAVGGAAGENQYIALTGVGEQVSSYLTYAKLAKNYKP.A...
sseJ_           WTEFLSSPHFL...........GKEMLNFAEGGSTS.ASYSCFNCIGDFVSNTD...RQVASYTPSH...
consensus>70    W.#%......L..............$.N.A.GG......Y.........!S...................

                220       230       240       250       260       270
BCT80_RS20435   KGDMVVLALSANDYMTFDKTDVEKVVNCYEKEIIKMVVSKGVKYILVTGIPDLSITAHAHKECD..RDRA
VPA0226         .NTLFTLEFGLNDFMNYNR.GVPEVKADYAEALIR.LTDAGAKNFMLMTLPDATKAPQFK.YSTQEEI.D
sseJ_           .QDLAIFLLGANDYMTLHKDNVIMVVEQQIDDIEK.IISGGVNNVLVMGIPDLSLTPYGK.HSD..EK.R
consensus>70    ...$.......ND%M......V..V...............G....$....PD..................

                280       290       300       310       320       330       340
BCT80_RS20435   KMSGLSNQHNQFLKSKLKDLQERFFGDGVKIGFFDFSGQLDELVDMANQVGYDTTT.....EEQAGYIDL
VPA0226         KIRAKVLEMNEFIKAQAMYYKA....QGYNITLFDTHALFETLTSAPEEHGFVNASDPCLDINRSSSVDY
sseJ_           KLKDESIAHNALLKTNVEELKEKY..PQHKICYYETADAFKVIMEAASNIGYDTEN....PYTHHGYVHV
consensus>70    K........N...K................I..%#.............#.G%...............!..

                350       360       370       380       390       400
BCT80_RS20435   PRVFGLKGNSQPFDSAHRHVFHDEVHPSQEVHQMLASRM.CD...FIVSQFGRTSKMNPSE
VPA0226         MYTHALRSECA.ASGAEKFVFWDVTHPTTATHRYVAEKMLESSNNLAEYR..........F
sseJ_           PGAKDPQ.....LDICPQYVFNDLVHPTQEVHHCFAIML.ES...FIAHHYST.......E
consensus>70    ...................VF.D..HP....H...A..$......................
```

### SignalP (6.0) analysis of BCT80_RS20435

**BCT80_RS20435**
**Prediction:** Other

| Protein type | Other | Signal Peptide (Sec/SPI) | Lipoprotein signal peptide (Sec/SPII) | TAT signal peptide (Tat/SPI) | TAT Lipoprotein signal peptide (Tat/SPII) | Pilin-like signal peptide (Sec/SPIII) |
|---|---|---|---|---|---|---|
| Likelihood | 1.0001 | 0 | 0 | 0 | 0 | 0 |

**Download:** PNG / EPS / Tabular

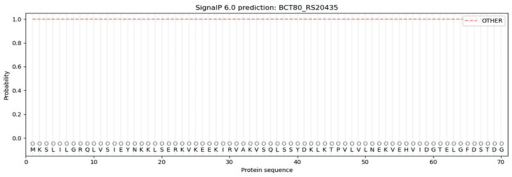


### SignalP (6.0) analysis of VPA0226

**Sequence**
**Prediction:** Signal Peptide (Sec/SPI)

Cleavage site between pos. 20 and 21. Probability 0.960276

| Protein type | Other | Signal Peptide (Sec/SPI) | Lipoprotein signal peptide (Sec/SPII) | TAT signal peptide (Tat/SPI) | TAT Lipoprotein signal peptide (Tat/SPII) | Pilin-like signal peptide (Sec/SPIII) |
|---|---|---|---|---|---|---|
| Likelihood | 0.0004 | 0.9989 | 0.0002 | 0.0002 | 0.0002 | 0.0002 |

**Download:** PNG / EPS / Tabular

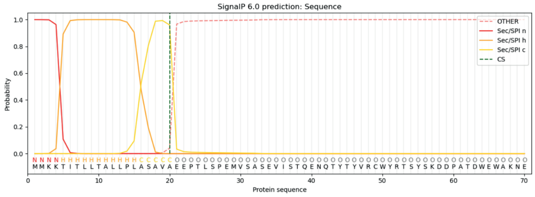
